# H5N1 2.3.4.4b HA E190D and Q226H mutations, picked up as minority variants in a patient, result in an inability to bind sialic acid

**DOI:** 10.64898/2026.03.06.710037

**Authors:** Eszter Kovács, María Ríos Carrasco, Mafalda F. Guerreiro Cabana, Robert P. de Vries

## Abstract

A human infection with clade 2.3.4.4b H5N1 influenza A virus in Canada revealed minority variants E190D and Q226H in the hemagglutinin (HA) receptor-binding site (RBS). Because mutations at positions 190 and 226 have been associated with altered receptor specificity in other influenza subtypes, we investigated their impact on receptor binding in H5 HA. Using a recombinant protein approach and an ELISA-based glycan-binding assay, we assessed binding to representative avian- and human-type sialylated glycans. Both single mutations and their combination resulted in a complete loss of detectable binding to the tested glycans. To evaluate whether this phenotype was background-dependent, Q226H was additionally introduced into two other H5 HA proteins, each representing a distinct clade. In both cases, the mutation similarly abolished receptor binding. These findings independently validate recent glycan microarray observations and demonstrate that the patient-derived E190D and Q226H substitutions severely impair receptor-binding capacity across multiple H5 backgrounds. Single mutations at key RBS residues in H5 often disrupt receptor binding rather than confer human-type receptor specificity, confirming complex mutational pathways required for adaptation to human-type receptors.

## Introduction

In November 2024, a 13-year-old girl was hospitalized in Canada due to a critical respiratory illness caused by influenza A virus (IAV) H5N1 (2.3.4.4b, A/British Columbia/PHL-2032/2024 [H5BC]). After sequencing, Q226H and E190D minority variants were detected in the receptor binding site (RBS) of the H5 hemagglutinin (HA) protein (1). Since minority mutations can also point to possible human adaptation, it is vital to analyze them for their effects on receptor-binding properties.

The HA receptor binding site (RBS) of H5 2.3.4.4b viruses retains several conserved structural elements found in other influenza A subtypes, including the canonical 130- and 220-loops and the 190-helix, as well as the conserved amino acid residues Y95, W153, H183, and Y195 that constitute the RBS(2). Although hallmark substitutions associated with the switch from avian-type (α2,3-linked sialic acids) to human-type (α2,6-linked sialic acids) receptor specificity have been well characterized for H1N1, H2N2, and H3N2 viruses (E190D/G225D and Q226L/G228S) (3), these simplified models do not directly apply to H5 subtypes (4). In H5 viruses, receptor-binding characteristics are shaped by a broader network of amino acid interactions within the RBS and adjacent regions, resulting in complex determinants of host adaptation (5). Recent analyses of clade 2.3.4.4 H5 viruses have shown that substitutions such as N224K, Q226L, and G228S can increase affinity for human-type receptors in vitro (6, 7).

In previous work, we assessed mutations that arose during human infection in classical H5 from the A/Indonesia/05/05 H5N1 strain and observed that most single-amino-acid mutations resulted in HA proteins with diminished receptor-binding properties (8). Additional mutations were necessary to gain human-type receptor specificity. Deep mutational scanning analyses of a 2.3.4.4b H5 protein revealed that mutations at positions 190 and 226 can broaden binding to a2,6-linked sialic acid on cells, yet 190D and 226H are not among them (9). Recently, it has been reported that the BC24 H5 HA protein lost all binding when the E190D and Q226H were introduced, while some fusion capacity was maintained (10). Here, using an ELISA-based complementary assay, we confirm this observation, which we deem important for open science, and that provides independent validation of these findings. Additionally, we introduced the Q226H in two different H5 antigenic backgrounds.

Thus, two human-infection-derived mutations result in a severe decrease in receptor binding, which might be a prerequisite for adaptation to human-type receptors.

## Results

### H5TX E190D, Q226H and E190D Q226H fail to bind common sialosides

We elected to introduce both E190D and Q226H as single and double mutants into the H5 HA of A/Texas/37/24 (H5TX), as it is a commonly employed human-derived HA. These proteins were tested for receptor-binding properties using a previously described glycan ELISA-based assay (7). Next to linear triLacNAc structures terminating with either α2,3- and α2,6-linked sialic acid, we added a symmetrical bi-antennary N-glycan with three LacNAc repeats terminating with α2,6-linked sialic acid on both arms, as this is a preferred human-type receptor (6, 11). Clearly, the introduction of either E190D or Q226H, as well as in combination, resulted in undetectable binding to commonly employed sialosides (Fig 1A). As a control for human-type receptor binding, we used a human H1N1 control (A/Kentucky/UR06-0258/07 H1N1), which efficiently bound both the α2,6-linked linear and biantennary N-linked glycan (12) (Fig 1B).

**Figure 1.**
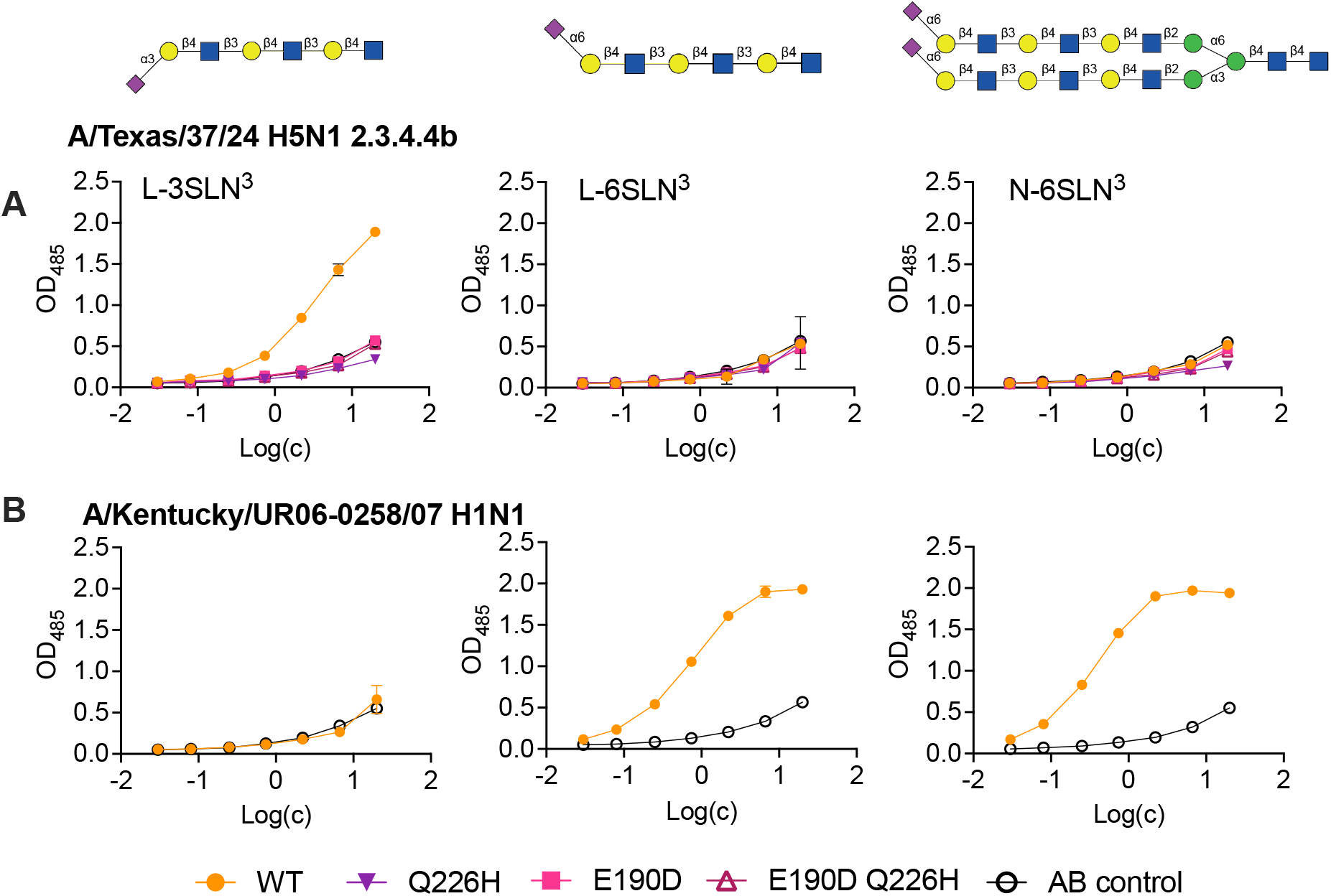
Direct receptor binding assay using biotinylated linear avian- and human-type receptors. Linear L-3SLN^3^ (left) L-6SLN^3^ (middle), and biantennary N-glycan N-6SLN^3^ coated streptavidin plates interrogated with A/Texas/37/24 and mutants E190D, Q226H and E190 Q226H mutants (**A**), and A/Kentucky/UR06-0258/07 H1N1 (**B**). The data are a representative of three biological independent assays performed in three technical replicates

### The Q226H mutation results in non-binders in two additional H5 backgrounds

Since the hallmark Q226L mutation can have different effects in different H5 backgrounds (7), we also analyzed the Q226H mutant in two additional H5 HA proteins. A/duck/France/161108h/16 (H5FR) is a representative of early 2.3.4.4b strains in Europe before the transmission to the American continent. This H5FR WT protein bound L-3SLN^3^, but the Q226H mutation abolished all binding (Figure 2A). A/black swan/Akita/1/16 2.3.4.4e H5 (H5AK) was selected as we previously observed binding to human-type receptors with a single Q226L mutation. The H5AK WT bound L-3SLN^3^; however, when Q226H was introduced, no binding to any of the glycans was detected (Figure 2A).

**Figure 2.**
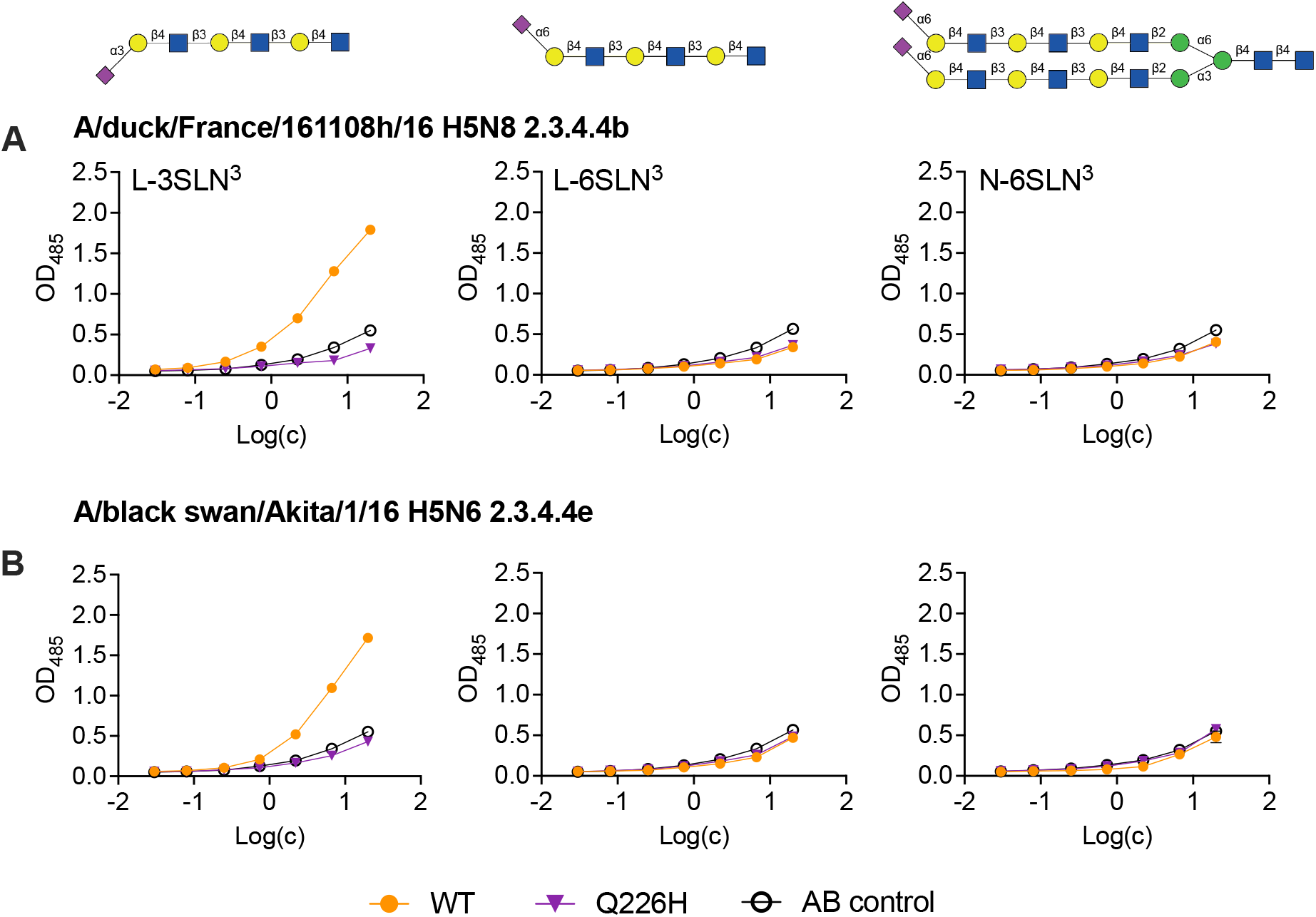
Direct receptor binding assay using biotinylated linear avian- and human-type receptors. Linear L-3SLN^3^ (left) L-6SLN^3^ (middle), and biantennary N-glycan N-6SLN^3^ coated streptavidin plates interrogated with A/duck/France/161108h/16 H5N8 and Q226H mutant (**A**), and A/black swan/Akita/1/16 and Q226H mutant (**B**). The data are a representative of three biological independent assays performed in three technical replicates

## Discussion

Using recombinant protein approaches, we demonstrate that the patient-identified E190D and Q226H mutations lead to H5 proteins with minimal sialic acid-binding properties. Our data is highly complementary to a recent preprint that also used glycan microarray approaches, in which no sialosides were bound by the H5BC E190D and/or Q226H mutants (10). We extend these analyses by introducing Q226H into several H5 protein backgrounds, all of which lead to severely diminished receptor-binding properties.

Previously, we generated several Q226 mutants in the classical A/Indonesia/05/05 H5N1 HA protein. Here, Q226A/R and L all lead to minimal binding on a simplified glycan microarray (8). Interestingly, these mutations were suggested to favor a2,6-linked sialic acid binding in a 2.3.4.4b HAs (9). A similar discrepancy is observed for E190A, although comparing receptor phenotypes of antigenic clade 2.1 with those of 2.3.4.4b HA is not accurate, as there are now multiple studies describing the uniqueness of the latter’s sialic acid-binding interactions (13-16).

Conclusively, single minority mutations in key sites in the RBS of H5 often lead to minimal receptor binding, as exemplified by E190D and Q226H. Our work complements and validates an earlier preprint (10), and confirms it is vital to analyze such mutants in different H5 backgrounds.

## Materials and methods

### Expression and purification of hemagglutinin

Codon-optimized HA-encoding cDNAs of different HA strains were cloned into the pCD5 expression vector as described previously (17). The pCD5 expression vector was modified to clone the HA-encoding cDNAs in frame with DNA sequences coding for a secretion signal sequence, an mOrange2 protein placed N-terminally, a GCN4 trimerization domain (RMKQIEDKIEEIESKQKKIENEIARIKK), and the Twin-Strep (WSHPQFEKGGGSGGGSWSHPQFEK; IBA, Germany) (17-19). The final constructs were transfected and purified from HEK293S GnTI(−) cells as described previously (17). In short, a mixture of DNA and polyethyleneimine I (PEI) in a 1:8 ratio (µg DNA: µg PEI) was added to the cells. The transfection mix was replaced by 293 SFM II suspension medium (Invitrogen, 11686029, supplemented with glucose 2.0 gram/L, sodium bicarbonate 3.6 gram/L, primatone 3.0 gram/L (Kerry), 1% glutaMAX (Gibco), 1.5% DMSO, and 2mM valproic acid after 6 hours. Five days post-transfection, supernatants were harvested, and proteins were purified using Strep-tactin Sepharose beads (IBA Life Sciences) as previously described (20).

### Glycan ELISA

Nunc MaxiSorp 96-well plates (Invitrogen) were coated with 50 μL of 5 μg/mL streptavidin (Westburg) in PBS overnight at 4°C. After blocking with 300 μL of 1% BSA in PBS-T for 3 hours at RT, streptavidin-coated plates were coated with 50 μL of 50 nM biotinylated glycans overnight at 4°C. After blocking with 300 μL of 1% BSA in PBS-T for 3 hours at room temperature. HAs at 20 μg/mL were precomplexed with strepmab and goat-*anti*-human antibodies (Invitrogen (#31410)) in a 1:0.65:0.325 molar ratio on ice for 30 min. Proteins were added to the plates, diluted serially 1:2, and incubated for 90 minutes at RT. After washing, the reaction was developed using an OPD solution and stopped with 2.5 M H_2_SO_4_ after 5 min. Absorbance was measured at 485nm (Polar Star Omega, BMG Labtech), and raw data were analyzed using GraphPad Prism.

## Acknowledgements

This research was made possible by funding from ICRAD, an ERA-NET co-funded under the European Union’s Horizon 2020 research and innovation program. (https://ec.europa.eu/programmes/horizon2020/en), under Grant Agreement n°862605 (Flu-Switch) to RPdV. MRC and RPdV are supported by an NWO-M2 (OCENW.M20.106). We would like to thank Dirk Eggink for the motivation.

